# Senolytic Intervention Improves Cognition, Metabolism, and Adiposity in Female APP^NL-F/NL-F^ Mice

**DOI:** 10.1101/2023.12.12.571277

**Authors:** Yimin Fang, Mackenzie R. Peck, Kathleen Quinn, Jenelle E. Chapman, David Medina, Samuel A. McFadden, Andrzej Bartke, Kevin N. Hascup, Erin R. Hascup

**Affiliations:** Department of Neurology, Dale and Deborah Smith Center for Alzheimer’s Research and Treatment, Neuroscience Institute, Southern Illinois University School of Medicine, Springfield, IL 62702, USA; Department of Internal Medicine, Southern Illinois University School of Medicine, Springfield, IL 62702, USA; Department of Medical Microbiology, Immunology and Cell Biology, Southern Illinois University School of Medicine, Springfield, IL 62702, USA; Department of Pharmacology, Southern Illinois University School of Medicine, Springfield, IL 62702, USA

**Author notes:** Corresponding Author: Erin R. Hascup, Department of Neurology, Dale and Deborah Smith Center for Alzheimer’s Research and Treatment, Southern Illinois University School of Medicine, Springfield, IL 62794-9268, USA.

**Keywords:** dasatinib and quercetin, fisetin, cell senescence, cognition, amyloid plaques, Alzheimer’s disease

## Abstract

Senescent cells accumulate throughout the body and brain contributing to unhealthy aging and Alzheimer’s disease (AD). The APP^NL-F/NL-F^ amyloidogenic AD mouse model exhibits increased markers of senescent cells and the senescence-associated secretory phenotype (SASP) in visceral white adipose tissue before plaque accumulation and cognitive decline. We hypothesized that senolytic intervention would alleviate cellular senescence thereby improving spatial memory in APP^NL-F/NL-F^ mice. Thus, four month old male and female APP^NL-F/NL-F^ mice were treated monthly with vehicle, 5 mg/kg Dasatinib + 50 mg/kg Quercetin, or 100 mg/kg Fisetin. Blood glucose levels, energy metabolism, spatial memory, amyloid burden, and senescent cell markers were assayed. Dasatinib + Quercetin treatment in female APP^NL-F/NL-F^ mice increased oxygen consumption and energy expenditure resulting in decreased body mass. White adipose tissue mass was decreased along with senescence markers, SASP, blood glucose, and plasma insulin and triglycerides. Hippocampal senescence markers and SASP were reduced along with soluble and insoluble amyloid-β (Aβ)_42_ and senescence associated-β-gal activity leading to improved spatial memory. Fisetin had negligible effects on these measures in female APP^NL-F/NL-F^ mice while neither senolytic intervention altered these parameters in the male mice. Considering women have a greater risk of dementia, identifying senotherapeutics appropriate for sex and disease stage is necessary for personalized medicine.

## Introduction

The most prevalent form of dementia, Alzheimer’s disease (AD), is an age-related neurodegenerative disorder that is considered the 6^th^ leading cause of death in the United States. Nearly 6 million individuals living in the United States are estimated to have an AD diagnosis, and this number is expected to double every five years. The pathological hallmarks of AD are senile plaques composed of aggregated Aβ and neurofibrillary tangles formed from hyperphosphorylated tau protein. Approved therapies are limited, with those targeting cholinergic and glutamatergic signaling only providing symptomatic relief. Recent anti-amyloid antibodies show modest abatement of cognitive decline, but have edema and microhemorrhage risks [1]. This has prompted the field to continue searching for alternative therapeutic targets.

Numerous research efforts have explored the pathophysiology and treatment of AD. Separate gene mutations causing familial AD and preventing sporadic AD, gave rise to the amyloid cascade hypothesis [2] that has become a centralized dogma underlying AD research for several decades [3]. This has led to an incomplete understanding of causal pathways and slowed progress in the discovery of novel treatments [4].

Currently, AD is considered a multifactorial disease involving tau hyperphosphorylation, Aβ accumulation, inflammation, mitochondrial dysfunction, oxidative stress, and dysregulated cellular communication culminating in neuronal loss [5]. Cellular senescence, as a fundamental contributor to both aging and AD [6–8], is defined as cell cycle arrest governed by complex interactions across p21 and p16 tumor suppressive pathways [9]. Senescent cells accumulate in nearly every tissue type of the body including the brain [10]. Their accumulation drives the production of senescent markers including p21, p16, and SASP [8, 11], which is involved in the expression of inflammatory cytokines, chemokines, growth factors, and matrix remodeling factors [12]. Over time the SASP profile can affect neighboring cells leading to a vicious cycle of cellular dysfunction.

Genetic deletion of senescent cells from the brain or treatment with senolytic drugs such as navitoclax [13], D+Q [14, 15], or Q [16, 17] has led to functional improvements in several different tau and amyloid AD models. The majority of these studies were conducted at later disease stages when pathology was pronounced. We hypothesized senolytics could be used as a primary prevention against AD. To test our hypothesis, we utilized a knock-in mouse model of AD (APP^NL-F/NL-F^) carrying the Swedish (NL) and Beyreuther/Iberian (F) amyloid precursor protein (APP) mutations [18, 19] that slowly develop plaque pathology starting at six months of age. From 4 to 13 months of age, male and female APP^NL-F/NL-F^ mice were treated monthly with either 5 mg/kg D + 50 mg/kg Q or 100 mg/kg Fisetin similar to our previous protocols [20]. The results revealed that treatment with D+Q improved cognitive function accompanied with reduced senolytic markers in the brain and adipose tissue and other physiological parameters in female APP^NL-F/NL-F^ mice.

## Methods

### Chemicals

Unless otherwise noted, all chemicals were obtained from Sigma-Aldrich (St. Louis, MO) including Quercetin (Cat# RHR1488). Fisetin was purchased from Selleckchem (Houston, TX; Cat #S2298), and Dasatinib from LC laboratories (Woburn, MA; Cat# D-3307).

### Animals

APP^NL-F/NL-F^ mice on a C57BL/6 background (RRID: IMSR_RBRC06343) were obtained from Riken (Japan). C57BL/6 mice were originally obtained from The Jackson Laboratory (Bar Harbor, ME) and used to maintain breeding colonies of both genotypes. Protocols for animal use were approved by the *Institutional Animal Care and Use Committee* at Southern Illinois University School of Medicine (Protocol number: 2022-055), which is accredited by the Association for Assessment and Accreditation of Laboratory Animal Care. All studies were conducted in accordance with the United States Public Health Service’s Policy on Humane Care and Use of Laboratory Animals. Mice were group housed on a 12:12 h light-dark cycle, with food (Chow 5001 with 23.4% protein, 5% fat, and 5.8% crude fiber; LabDiet PMI Feeds) and water available *ad libitum*. The majority of experiments were conducted during the light phase except for indirect calorimetry where data was obtained for a 24 hr cycle. Genotypes were confirmed by collecting a 5 mm tail tip for DNA analysis by TransnetYX, Inc (Cordova, TN).

### Senolytic treatment

Senotherapeutic concentrations and dosing strategy were based on previous publications [20–22]. APP^NL-F/NL-F^ mice were dosed with 100 mg/kg of Fisetin, 5 mg/kg of D + 50 mg/kg of Q, or vehicle (2% DMSO in canola oil) by oropharyngeal administration similar to our previous studies [20]. The treatments were given once per month from 4-13 months of age. All assays, including tissue collection, were conducted one week after monthly senotherapeutic dosing. The dosing strategy was based on previous literature showing that markers of cell senescence are still reduced after a four week off-treatment period [8]. Since senescent cell accumulation occurs over weeks, maintaining an effective concentration dosage is not required. Rather, intermittent dosing may be preferable thereby avoiding adverse effects.

### Body weight and chow consumption monitoring

Mouse body weight and chow consumption were monitored weekly with the former used for senolytic dosing. Chow consumption was determined by dividing the amount eaten by the total body weight (g) of mice in each cage. Cages were averaged across groups for each month of treatment.

### Body composition and tissue procurement

A week following the final senotherapeutic treatment at 13 months of age, body weight was recorded and mice were sedated using isoflurane. A cardiac puncture was used to collect the plasma. Mice were rapidly decapitated and the peripheral tissues (liver, heart, kidney, pancreas, spleen, white, and brown adipose tissue [WAT; BAT]) were weighed to calculate their contribution to total body weight. Tissue was immediately flash frozen and stored at -80°C until processing as previously described [20].

### Glucose tolerance test (GTT) and insulin tolerance test (ITT)

Glucose metabolism was assessed via GTT and ITT after six treatments at the animal age of nine months as previously described [20]. Fifteen-hour-fasted mice underwent GTT by intraperitoneal (i.p.) injection with 1 g glucose per kg of body weight (BW). Blood glucose levels were measured at 0, 15, 30, 45, 60, and 120 min with a PRESTO glucometer (AgaMatrix). For ITT, nonfasted mice were injected i.p. with 1 IU porcine insulin from sigma (St. Louis, MO) (Cat# I5523) per kg of BW. Blood glucose levels were measured at 0, 15, 30, 45, 60, and 120 min. The data for GTT and ITT is presented as absolute value.

### Indirect calorimetry

Energy metabolism was measured by indirect calorimetry at the animal age of twelve months using an AccuScan Metabolic System (AccuScan Instruments). In this system, mice are housed individually in metabolic chambers with *ad libitum* access to food and water. After a twenty-four-hour acclimation period, the oxygen consumption (VO_2_), carbon dioxide produced (VCO_2_), energy expenditure (EE), and respiratory quotient (RQ) measurements were collected every ten minutes per animal and averaged for each hour.

### Morris water maze (MWM) training and probe challenge

The MWM was used to assess spatial learning and memory recall, and performed as previously described [23]. Mice were trained to utilize visual cues placed around the room to repeatedly swim to a static, hidden escape platform (submerged one cm below the opaque water surface), regardless of starting quadrant. The MWM paradigm consisted of 5 consecutive training days with three, 90 s trials/day and a minimum inter-trial-interval of 20 min. Starting quadrant was randomized for each trial. After two days without testing, the escape platform was removed and all mice entered the pool of water from the same starting position for a single 60 s probe challenge to test long-term memory recall. The ANY-maze video tracking system (Stoelting Co., Wood Dale, IL) was used to record mouse navigation during the training and probe challenge. The three trials for each training day were averaged for each mouse.

### Soluble Aβ_42_ Determination

The hippocampus from one hemisphere was dissected and stored at -80°C until tissue processing. Soluble Aβ_42_ concentrations were determined using the Human / Rat β amyloid ELISA kits (WAKO Chemicals; Cat: 292-64501) according to the manufacturer recommended protocols.

### Hippocampal amyloid plaque and acidic senescence-associated-β-galactosidase (SA-β-gal) staining and quantification

Immediately following dissection, hemibrains were fixed in 4% paraformaldehyde then transferred into 30% sucrose in 0.1M phosphate buffer prior to cryosectioning [24]. Twenty micron coronal sections through the hippocampus were obtained and every sixth serial section was stained for senescent cells and amyloid plaques. Free floating slices were stained for SA-β-gal per the manufacturer’s protocol (Cell Signaling Technology Danvers, MA, Cat#9860). Each slice was rinsed with 1X PBS then transferred to 1X Fixative Solution for 10-15 minutes followed by rinsing twice with 1X PBS. Slices were placed in β-gal staining solution (pH=6.0) and incubated for ∼18 hrs at 37°C. Following overnight incubation, slices were rinsed in 1X PBS, then mounted onto subbed microslides and allowed to dry before staining for amyloid plaques with 1% Congo Red (1 hr) according to the methods of Rajamohamedsait and Sigurdsson, 2012 [25]. After staining, slides were dehydrated using increasing concentrations of ethanol followed by submersion in CitriSolv for 5-10 min then immediately cover slipped using Permount (Fisher Scientific, Cat# SP15-500). Samples were imaged in brightfield at 4x and 40x magnification using a Keyence BZ-X810 All-in-One Fluorescence Microscope and accompanying analysis software (Keyence Corp.). The hippocampus was manually outlined and regions of interest were extracted by hue for each stain using the software’s Hybrid Cell Count function that determined plaque number and size, as well as the area of the hippocampus, plaques, and SA-β-gal stained regions. Senescence and plaque burden were calculated as a percentage of SA-β-gal-stained area within the hippocampus for each sample [26, 27]. Verification and quantification of amyloid plaques was conducted according to our previous protocols [28].Four hippocampal slices were averaged to obtain an individual sample for each mouse.

### Assessment of blood chemistry

Plasma was collected from animals anesthetized with isoflurane by cardiac puncture at the time of sacrifice. The blood was mixed with EDTA, followed by centrifugation at 10,000 g for 15 min at 4°C for plasma collection. Plasma was frozen at -80°C until determination of insulin and adiponectin (Crystal Chem, Elk Grove Village, IL; Cat# 90080 and 80569) by ELISA. Triglycerides or cholesterol (Pointe Scientific, Canton, MI; Cat# T7532-120 and C7510-120) was measured using the manufacturers’ recommended protocols. Plasma cytokine levels were assayed using a multiplex immunoassay (V-PLEX Custom Mouse Assay Kit) on a MESO QuickPlex SQ 120 with accompanying software (Meso Scale Diagnostics, LLC).

### RT–PCR

mRNA expression was analyzed by quantitative RT–PCR as previously described using cDNA synthesis kits and SYBR green mix from Bio-Rad (Cat# 1708897 and 1725121) performed with the StepOne Real-Time PCR System (Thermo Fisher Scientific) or the QuantStudio PCR System (Applied Biosystems). RNA was extracted using RNeasy mini kits or RNeasy Lipid Tissue Mini Kits (Qiagen) following the manufacturer’s instructions. Relative expression was calculated as previously described [29] and primers were purchased from Integrated DNA Technologies. The housekeeping genes glyceraldehyde 3-phosphate dehydrogenase (GAPDH) was used for adipose tissue while ubiquitin conjugating enzyme E2 D2 (UBE2D2) was used for the hippocampus. A list of forward and reverse primers is shown in Table 1.

**Table 1.**
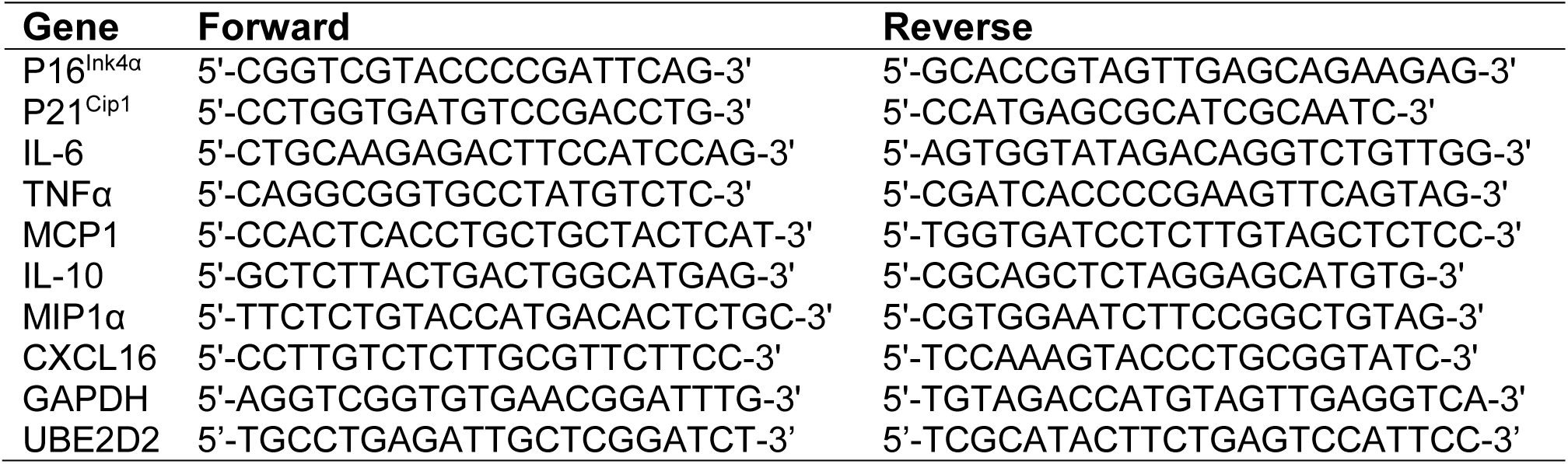
A list of forward and reverse primers used in these studies.

### Statistical analysis

Statistical analyses were conducted using a two-way ANOVA to test for significance of sex (males × females), treatment (Vehicle × Fisetin or D+Q), and interaction effects (sex x treatment). Differences between vehicle and senolytic treatments were determined with unpaired two-tailed Student’s t tests with significance defined as p < 0.05. Data are presented as means ± SEM. All statistical analyses and graphs were completed using Prism 10 (GraphPad Inc, La Jolla, CA, USA).

## Results

### APP^NL-F/NL-F^ mice have increased senescent cell accumulation and SASP in visceral white adipose tissue

Relative expression of senescent and SASP markers were assayed in visceral WAT (vWAT) from a separate cohort of four month old C57BL/6 and APP^NL-F/NL-F^ mice. Four month old female APP^NL-F/NL-F^ mice had increased expression of p16^Ink4α^, p21^Cip1^, IL-6, TNFα, MCP1, IL-10, MIP1α, and CXCL16 compared to age-matched female C57BL/6 mice (Online Resource 1A-H). p21^Cip1^, IL-6, TNFα, IL-10, and MIP1α were increased in male APP^NL-F/NL-F^ mice compared to four months old male C57BL/6 controls (Online Resource 1B-D, F-G). This data indicated knock-in of APP mutations increased senescent and SASP markers by four months of age, which made this an ideal therapeutic time point to administer senolytic treatment. Also, APP^NL-F/NL-F^ mice at this age do not have overt signs onset of cognitive deficits making it an optimal period for primary prevention treatment to determine disease-modifying outcomes.

### D+Q treatment decreases body weight via reduced adiposity in female APP^NL-F/NL-F^ mice

Metabolic deficits leading to increased oxidative stress has been observed in AD patients and transgenic mouse models prior to plaque and tangle development [30–32]. In the APP^NL-F/NL-F^ mice, Aβ deposition begins at 6 months of age with cognitive impairments occurring by 12-18 months [18]. To examine the metabolic changes associated with senolytic administration, body composition, glucose utilization, and energy metabolism were measured. Starting body weight was similar for each treatment group within a sex and monitored weekly thereafter (Online Resource 2A). Chow consumption was monitored weekly and no differences were observed throughout the treatment regimen (Online Resource 2B-C). Immediately before euthanization, mice were weighed to determine body composition. When compared to control treatment, D+Q decreased body weight while an increase was observed in Fisetin-treated female APP^NL-F/NL-F^ mice. Male APP^NL-F/NL-F^ mice had no change in body weight with either senolytic treatment (Fig. 1A). We examined body composition of BAT and WAT depots to determine changes from senolytic administration. The percentages of total WAT consisting of subcutaneous WAT, vWAT, and retroperitoneal WAT was reduced in female APP^NL-F/NL-F^ after D+Q treatment, but increased with Fisetin. Neither senolytic treatment effected WAT in males (Fig. 1B-E). Fisetin increased interscapular BAT in female APP^NL-F/NL-F^ mice but no changes were observed in the other groups (Fig. 1F). Body weight treatment effects were largely influenced by the changes in adiposity. The observed body composition differences could impact circulating lipids promoting dyslipidemia which increases the risk for developing cognitive impairments. Plasma triglyceride levels were decreased solely in females undergoing D+Q treatment but no changes were found in the other groups compared with control treatment (Fig. 1G). Cholesterol levels in plasma did not change with either of the treatment conditions in both the male and female mice (Fig.1H). In summary, D+Q treatment reduced WAT, body weight, and circulating triglycerides while Fisetin treatment increased WAT, BAT, and body weight in female APP^NL-F/NL-F^ mice. Neither D+Q nor Fisetin affected these measures in male APP^NL-F/NL-F^ mice.

**Figure 1.**
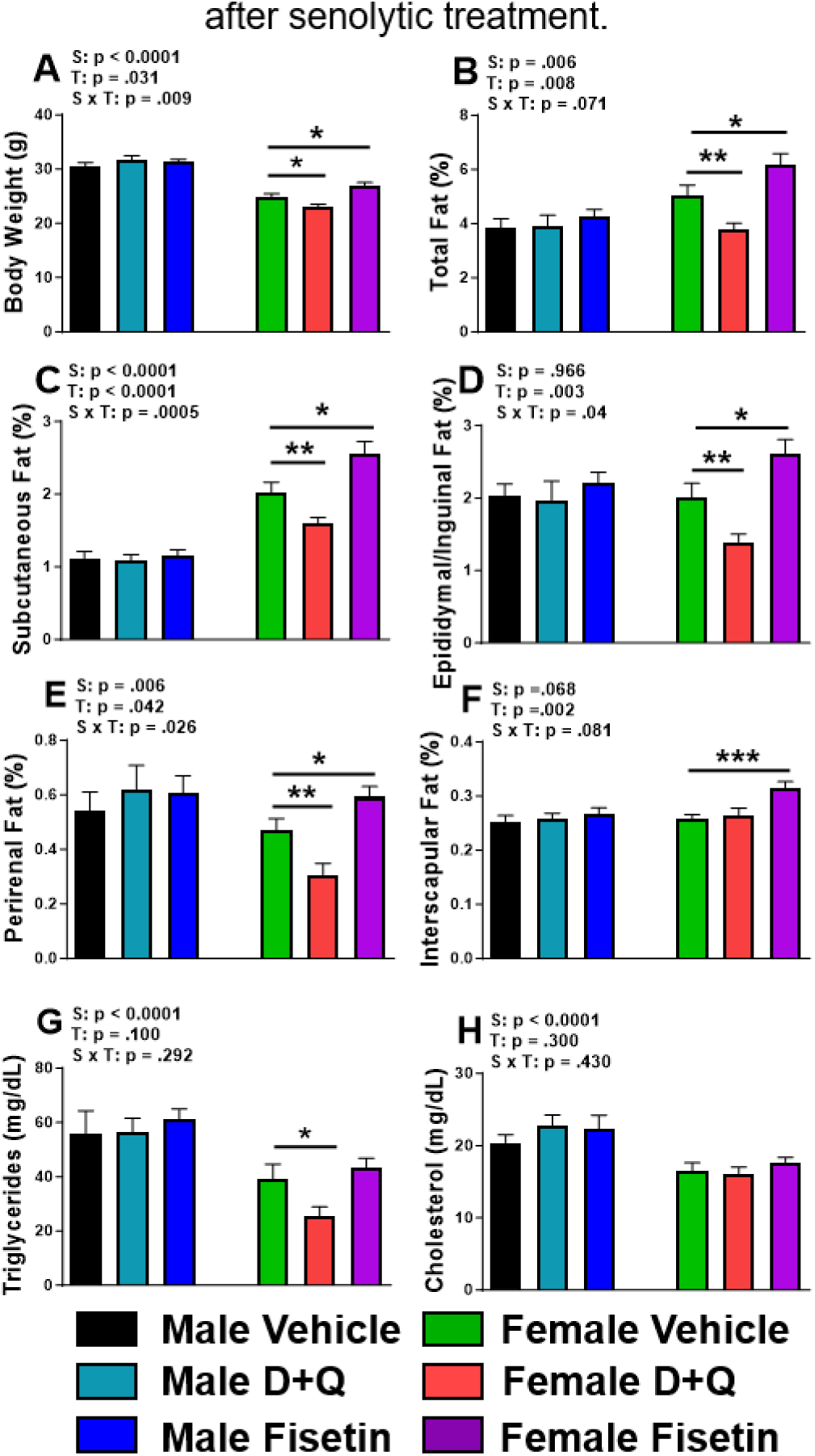
Senolytic intervention reduced adiposity and circulating lipid levels in female APP^NL-F/NL-F^ mice. Mouse body weight (A) along with percentages of total (B), WAT depots (C-E), and BAT (F) in relation to body weight. Plasma triglyceride and cholesterol levels were determined by enzymatic determination (G-H). All tissue was collected at the time of euthanization (14 months of age). Data are represented as means ± SEM (n=15-21). Results of a two factorial analysis are shown above each bar graph for the Sex (S) and Treatment (T) categorial variables and their interaction (S x T). *p<0.05, **p<0.01, ***p<0.001 based on a two-tailed Student’s *t* test.

### Effects of senolytic drug treatments on glucose metabolism in APP^NL-F/NL-F^ mice

Adipose tissue modulates glucose homeostasis, so we assayed the effects of senolytic intervention on blood glucose levels in APP^NL-F/NL-F^ mice. A GTT for glucose clearance and ITT for insulin sensitivity were performed. GTT and ITT results were unchanged regardless of sex or senolytic treatment compared with control groups (Fig. 2A-F). In female APP^NL-F/NL-F^ mice, D+Q and Fisetin treatment reduced fasting glucose, while D+Q treatment lowered fed glucose levels and plasma insulin (Fig. 2G-I). Fisetin and D+Q increased male APP^NL-F/NL-F^ plasma insulin levels (Fig 2I). D+Q treatment drastically reduced circulating adiponectin levels in male APP^NL-F/NL-F^ mice (Fig. 2J). Despite these changes to plasma insulin and adiponectin in male APP^NL-F/NL-F^ mice, no corresponding blood glucose changes were observed with the GTT or ITT (Fig. 2A,C,E-H).

**Figure 2.**
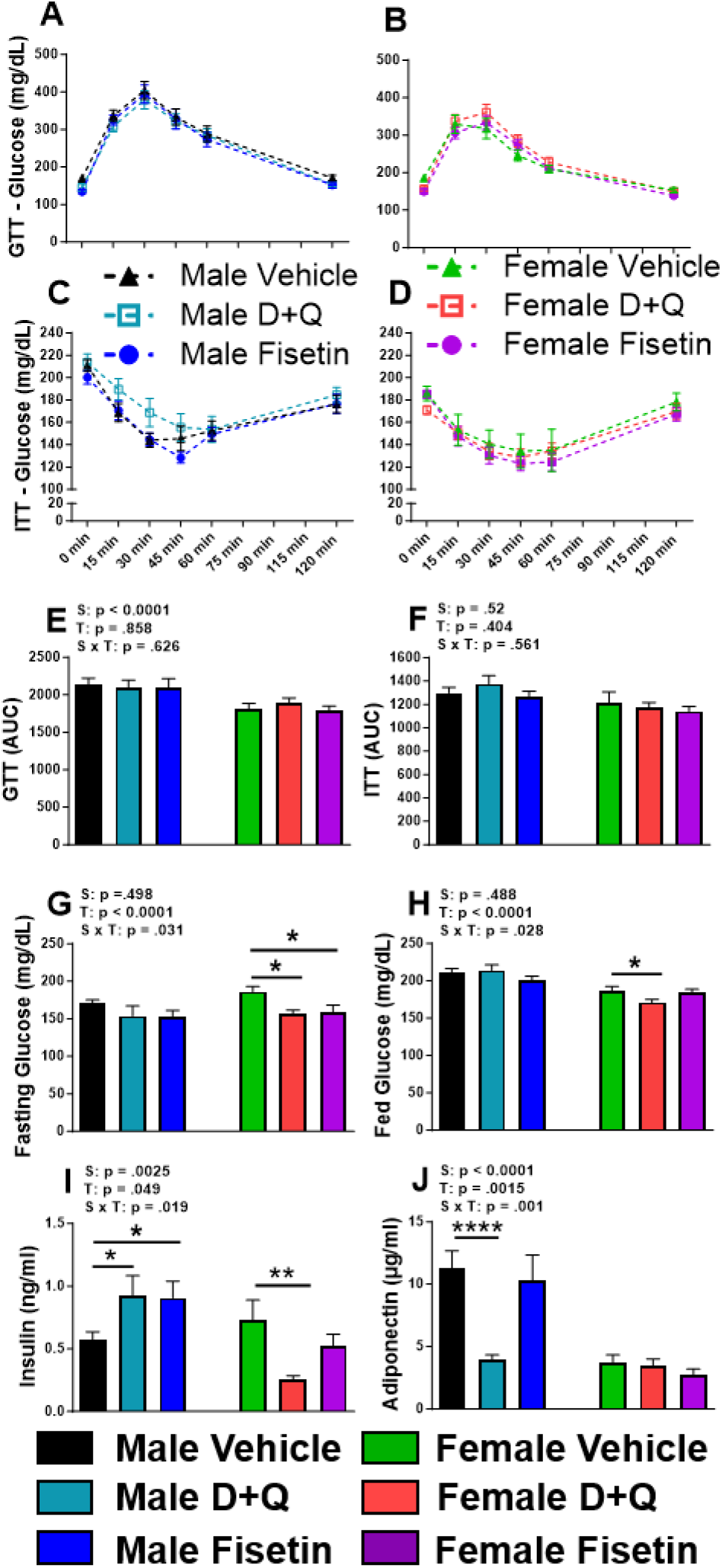
Blood glucose and plasma insulin are improved after senolytic treatment in female APP^NL-F/NL-F^ mice. Glucose tolerance test (GTT) and insulin tolerance test (ITT), and their respective area under the curve (AUC) after five treatments (A-F). Fasting (G) and fed (H) blood glucose levels were determined from time=0 during the GTT and ITT, respectively. Circulating insulin and adiponectin levels were determined by ELISA from plasma collected at the time of euthanization (10 treatments). Data are represented as means ± SEM (n=13-23). Results of a two factorial analysis are shown above each bar graph for the Sex (S) and Treatment (T) categorial variables and their interaction (S x T). *p<0.05, **p<0.01, ****p<0.0001 based on a two-tailed Student’s *t* test.

### Effects of senolytic drug treatments on energy expenditure in APP^NL-F/NL-F^ mice

Aside from being an important regulator of glucose homeostasis, adipose tissue is also a central metabolic organ in the regulation of energy homeostasis [33]. Since D+Q and Fisetin senolytic drug treatments altered adipose tissue composition (Fig. 1A-E), indirect calorimetry was used to assess senolytic treatment effects on energy metabolism in APP^NL-F/NL-F^ mice. Data revealed that D+Q treatment elevated VO_2_ (Fig. 3B-C) and EE (Fig. 3E-F) without affecting respiratory quotient RQ (Fig 3H-I) in the female APP^NL-F/NL-F^ mice, while Fisetin elicited no effects. In contrast, neither treatment altered the energy metabolism in male APP^NL-F/NL-F^ mice (Fig. 3A,C,D,F,G,I).

**Figure 3.**
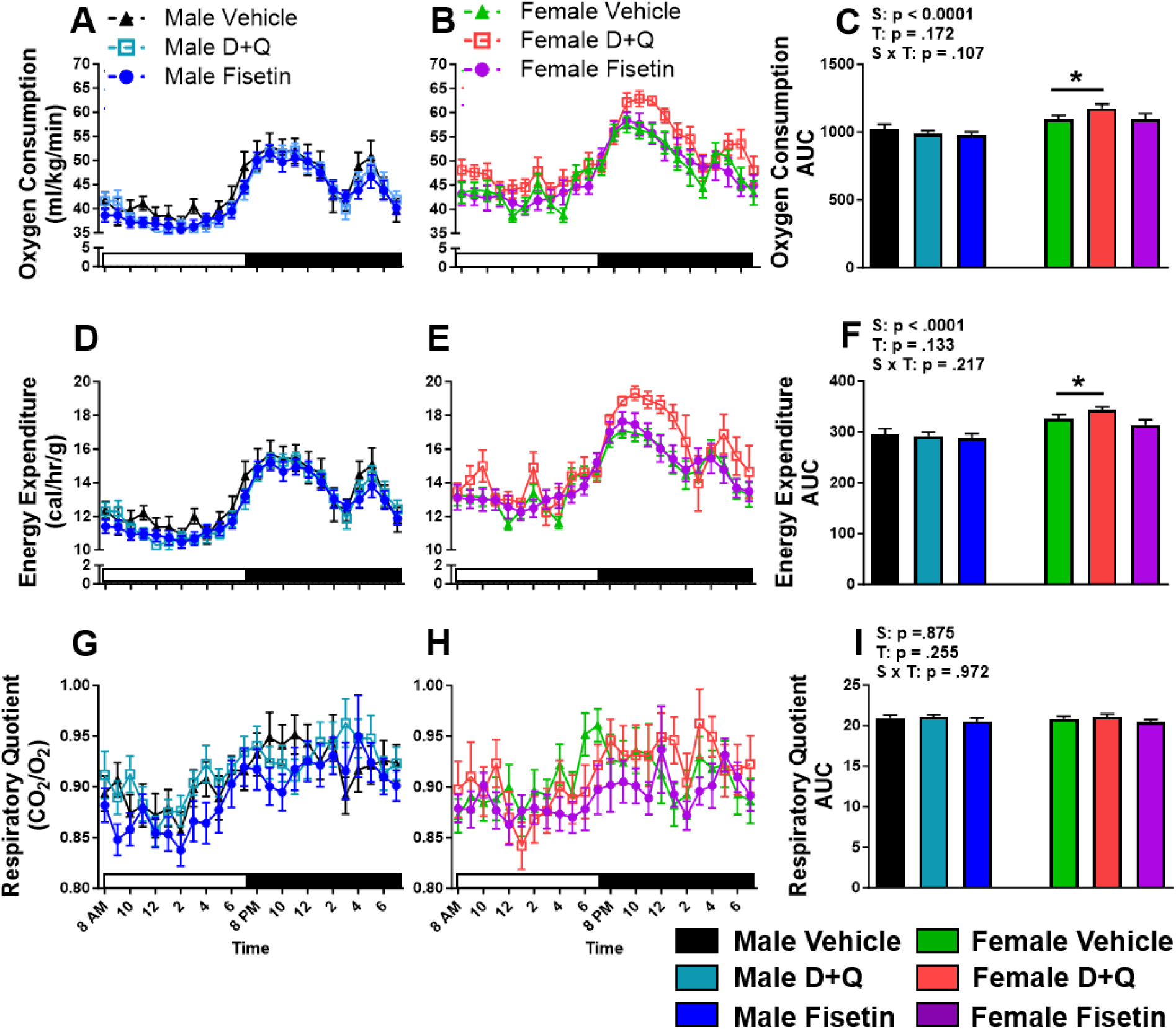
D+Q treatment improves oxygen consumption and energy expenditure in female APP^NL-F/NL-F^ mice. Twenty-four hour male and female oxygen consumption (A-C), energy expenditure (D-F), and respiratory quotient (G-I; VCO_2_/VO_2_) along with their corresponding AUC after 9 treatments were determined by indirect calorimetry. White and black bars along the abscissa indicates the hours of lights on or off, respectively. Data are represented as means ± SEM (n=15-23). Results of a two factorial analysis are shown above each bar graph for the Sex (S) and Treatment (T) categorial variables and their interaction (S x T). *p<0.05 based on a two-tailed Student’s *t* test.

### D+Q treatment reduces visceral white adipose tissue and hippocampal senolytic markers in female APP^NL-F/NL-F^ mice

As previously discussed, D+Q and Fisetin have been reported to reduce senescent cell burden in numerous models of aging and disease [8]. Adipose tissue, specifically vWAT, contains the highest burden of senescent cells [34] and has a stronger correlation with brain atrophy than age, BMI, hypertension, or type 2 diabetes mellitus [35]. Since D+Q and Fisetin treatment altered adiposity in female APP^NL-F/NL-F^ mice (albeit with contrasting effects), we further examined whether senescent cell markers and SASP were affected in vWAT. The results showed that D+Q treatment downregulated gene expression of p16^Ink4a^, p21^Cip1^, IL-6, TNFα, MCP1, and IL-10 within female APP^NL-F/NL-F^ mice (Fig. 4A-F). In contrast, D+Q treatment in male APP^NL-F/NL-F^ mice increased gene expression of p21^Cip1^, IL-6, TNFα, MCP1, IL-10, MIP1α, and CXCL16 (Fig. 4B-H) in vWAT. Fisetin treatment had no effect on vWAT genes in either sex of APP^NL-F/NL-F^ mice. Within the hippocampus, relative expression of p16^Ink4a^, p21^Cip1^, and TNFα (Fig 4I-K) were all decreased in D+Q treated female APP^NL-F/NL-F^ mice, similar to vWAT. D+Q treated males also had reduced TNFα hippocampal expression, while no effects were observed in either sex after Fisetin treatment. Overall, D+Q treatment reduced central and peripheral senescent and SASP makers in female APP^NL-F/NL-F^ mice.

**Figure 4.**
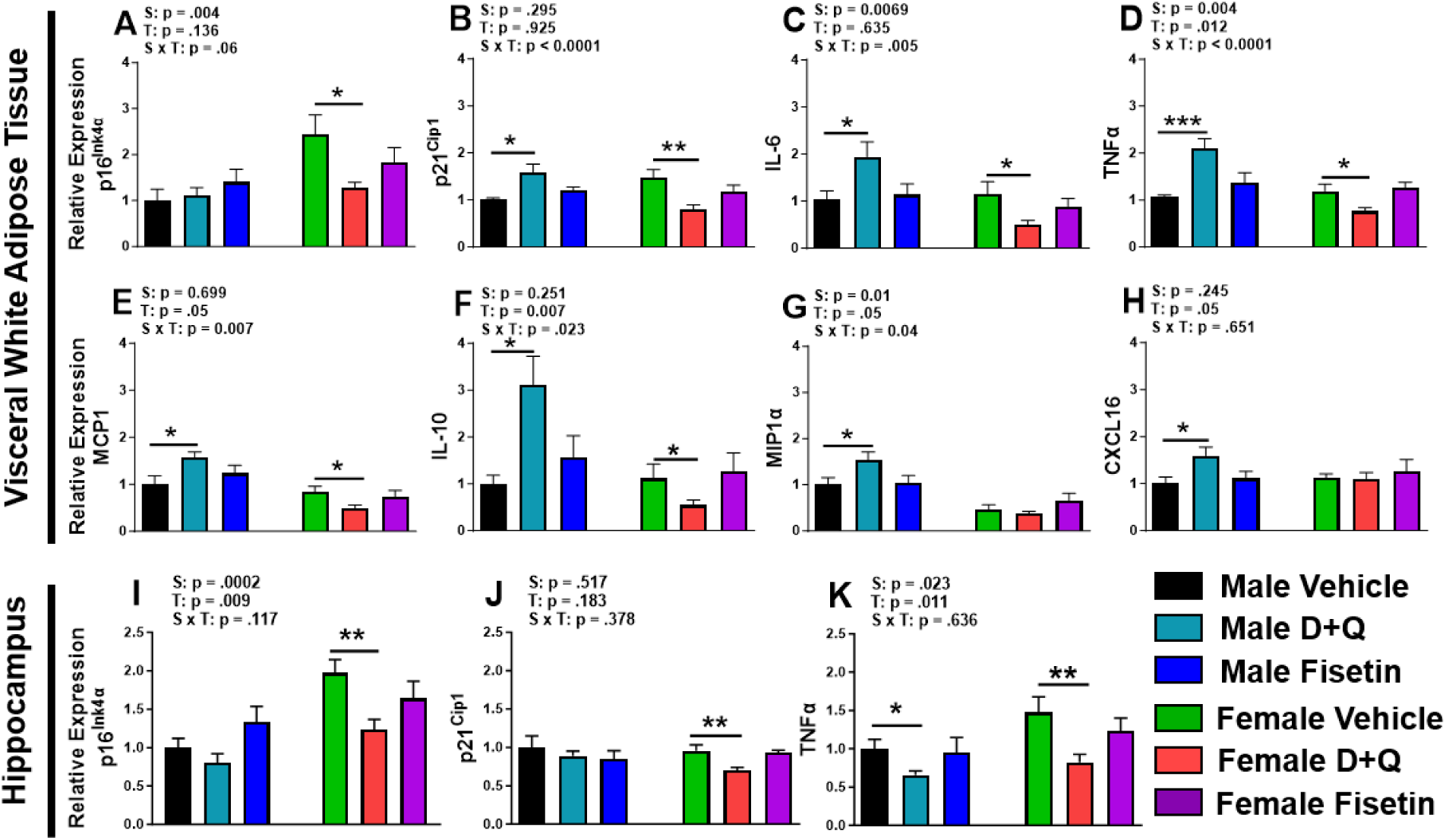
D+Q treatment altered senescent cell markers and SASP profile in a sexually dimorphic manner. Gene expression profiles in vWAT (A-H) and the hippocampus (I-K). Data are represented as means ± SEM (n=8-10). Results of a two factorial analysis are shown above each bar graph for the Sex (S) and Treatment (T) categorial variables and their interaction (S x T). *p<0.05, **p<0.01, ***p<0.001 based on a two-tailed Student’s *t* test.

### Senolytic treatment reduces plasma cytokine levels in female APP^NL-F/NL-F^ mice

Since senescent cell burden is related to secretion of SASP, we examined plasma cytokines by multiplex analysis (Online Resource 3A-E). D+Q treatment in female APP^NL-F/NL-F^ reduced plasma expression of IL-6, TNFα, and IL-1β, while fisetin treatment also reduced these three proinflammatory cytokines and IL-10. Despite elevated vWAT SASP expression in male APP^NL-F/NL-F^ mice, neither senolytic treatment altered plasma cytokine levels.

### D+Q treatment reduces hippocampal plaque burden in female APP^NL-F/NL-F^ mice

Higher senescence and SASP markers in adipose tissue has been linked to cognitive deficits and increased dementia risk [36–38]. Accordingly, it is important to know whether the senolytic drug treatments mitigate senescence and amyloid plaques in the brain of APP^NL-F/NL-F^ mice. SA-β-gal staining has been extensively used as a marker of cellular senescence *in vivo* in both whole-mount and cryosections [39]. To examine the treatment effects on cell senescence and amyloid plaques in the hippocampus, tissue was co-stained with SA-β-gal for senescence (blue) and congo red for amyloid plaques (red). The representative stained images (Fig. 5A) and quantified data showed that D+Q treatment in the female APP^NL-F/NL-F^ mice reduced the percentage of SA-β-gal staining (Fig. 5B) and the plaque number, size, density, and burden throughout the hippocampus (Fig. 5C-F). Since soluble Aβ_42_ is considered the neurotoxic species associated with AD development and its aggregation results in insoluble plaque deposition, we also assayed its hippocampal levels. Hippocampal soluble Aβ_42_ was reduced in D+Q treated female APP^NL-F/NL-F^ mice (Fig 5G). D+Q treatment in male APP^NL-F/NL-F^ mice as well as Fisetin treatment in both sexes did not drastically alter hippocampal soluble or insoluble amyloid levels (Fig. 5C-G). Furthermore, a sex effect was discovered with respect to males portraying a smaller plaque size than females within the control condition (Fig. 5D), which is consistent with our previous observations [28]. However, this sex effect did not carry over into any of the treatment groups.

**Figure 5.**
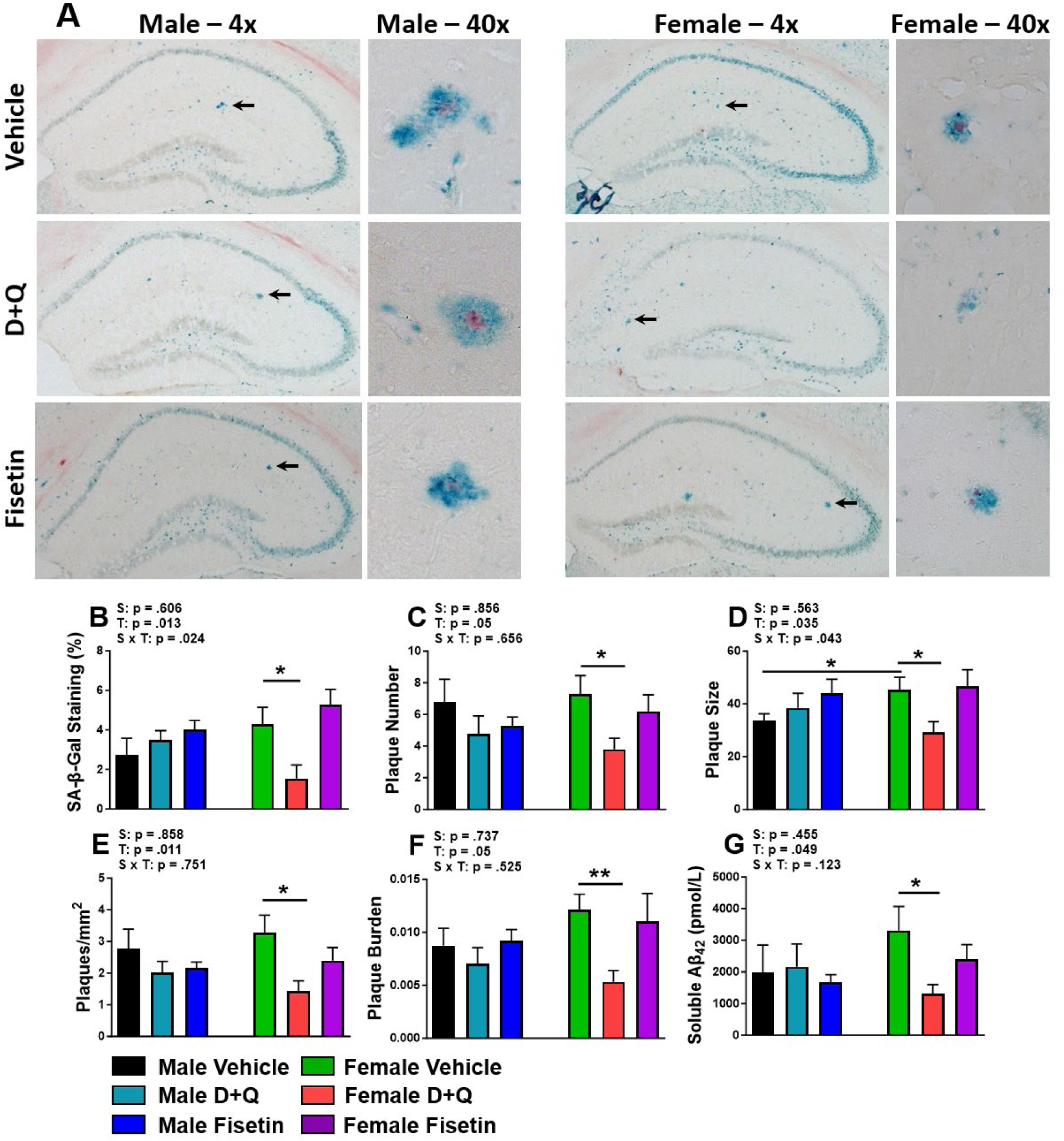
APP^NL-F/NL-F^ mice have reduced hippocampal senescent cell burden as well as insoluble and soluble Aβ_42_ after D+Q treatment. Representative 4x and 40x hippocampal images (A) of SA-β-gal staining (blue) and plaque accumulation (red) for male (left) and female (right) after 10 treatments of vehicle (top), D+Q (middle), or Fisetin (bottom). Arrows indicate magnified image area shown to the right of each whole hippocampal image. Percentage of SA-β-gal staining (B) along with insoluble (C-F) and soluble Aβ_42_ (G) levels are shown. Data are represented as means ± SEM (n=8-10). Results of a two factorial analysis are shown above each bar graph for the Sex (S) and Treatment (T) categorial variables and their interaction (S x T). *p<0.05, **p<0.01 based on a two-tailed Student’s t test.

### D+Q treatment improved spatial learning and memory in female APP^NL-F/NL-F^ mice

To examine whether senolytic treatment improves spatial learning and memory recall in APP^NL-F/NL-F^ mice, the MWM test was performed after 9 months of treatments at 13 months of age. The MWM data demonstrated that D+Q treated female APP^NL-F/NL-F^ mice improved spatial learning during the 5-day training period by reducing cumulative distance (Fig. 6B-C), decreasing corrected integrated path length (CIPL; Fig. 6E-F), and increasing path efficiency (Fig. 6H-I). D+Q treatment also improved spatial memory recall by significantly increasing platform and annulus 40 entries during the probe challenge (Fig. 6J-K). Fisetin treatment in female APP^NL-F/NL-F^ mice had minimal effects on spatial navigation, though an increase in path efficiency during the training days was observed (Fig. 6H-I). MWM spatial learning and memory recall was not altered in male APP^NL-F/NL-F^ mice receiving either D+Q or Fisetin (Fig. 4A,C,D,F,G,I-K). We did not observe sex or treatment effects with novel object recognition memory (data not shown). The improved spatial recognition learning and memory observed in female APP^NL-F/NL-F^ may be due to the reduced SASP profile as well as soluble and insoluble amyloid accumulation.

**Figure 6.**
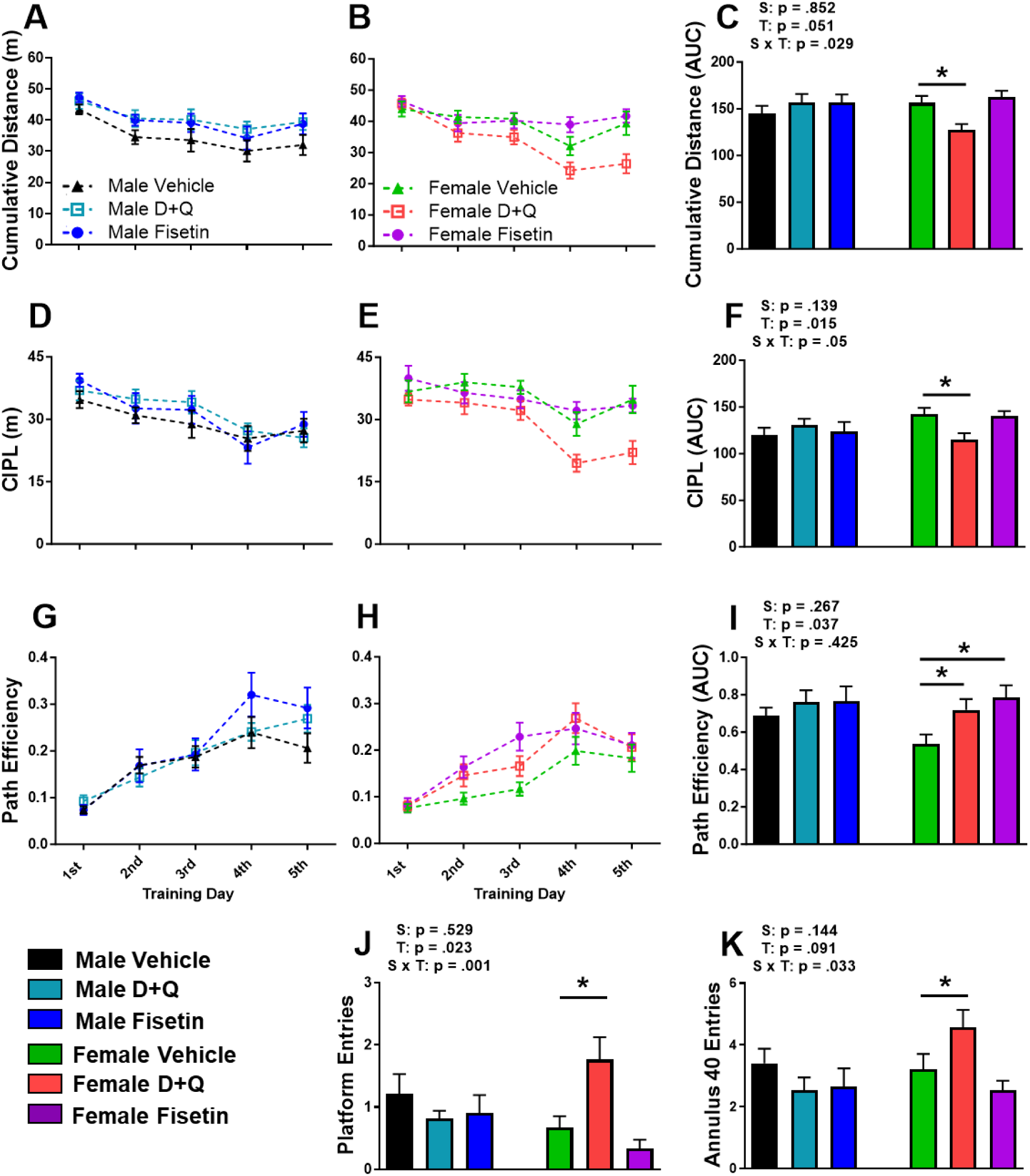
Spatial learning and memory recall was improved in female APP^NL-F/NL-F^ mice receiving D+Q treatment. An eight day MWM paradigm consisting of 5 training days followed by a single probe challenge 48 hrs after was used to assess spatial cognition after the eighth treatment (12 months of age). The cumulative distance (A-C), corrected integrated path length (CIPL; D-F), and path efficiency (G-I) along with their corresponding AUC are shown for each training day. The number of platform crosses into the former location of the hidden escape platform (J) and annulus 40 (K) are reported for the probe challenge. Data are represented as means ± SEM (n=20-25). Results of a two factorial analysis are shown above each bar graph for the Sex (S) and Treatment (T) categorial variables and their interaction (S x T). *p<0.05 based on a two-tailed Student’s *t* test.

## Discussion

Primary prevention using senolytics had sexually dimorphic effects in APP^NL-F/NL-F^ mice. D+Q treatment in female APP^NL-F/NL-F^ mice improved multiple peripheral and metabolic parameters including reduced body weight and adiposity, leading to improvements in triglyceride, insulin, and blood glucose levels as well as enhanced energy metabolism. D+Q treatment in female APP^NL-F/NL-F^ mice also decreased SASP markers in vWAT, plasma, and hippocampus. Furthermore, D+Q treatment in female APP^NL-F/NL-F^ mice decreased hippocampal soluble and insoluble Aβ_42_ and SA-β-gal staining. The culmination of these factors positively impacted spatial learning and memory in female APP^NL-F/NL-F^ mice. In male APP^NL-F/NL-F^ mice, D+Q treatment had an opposing effect whereby senescent cell markers and the SASP profile was increased while negatively impacting circulating insulin and adiponectin. Although these senescence and plasma profiles were worse, this did not alter overall body composition, blood glucose levels, energy metabolism, plaque burden, nor cognitive impairments beyond what was reported for vehicle treatment.

Fisetin treatment in male APP^NL-F/NL-F^ mice only negatively impacted plasma insulin levels. In the females receiving fisetin, we observed increased adiposity, unaffected hippocampal or vWAT senescent and SASP markers, but reduced circulating plasma cytokine levels. Opposing effects within the same sex were also observed between the two senolytic treatments. Expression of p21^Cip1^ and SASP vWAT markers were elevated in males receiving D+Q, unlike fisetin treatment. D+Q treatment in males reduced plasma adiponectin levels that would increase the proinflammatory profile of vWAT. In female APP^NL-F/NL-F^ mice, opposing senolytic treatment effects were predominantly observed for body weight and adipose tissue accumulation that we attributed to differences in serum triglyceride levels. Fisetin treatment also improved fasting blood glucose and several plasma proinflammatory cytokines in female

APP^NL-F/NL-F^ mice. Path efficiency during the MWM learning sessions was observed in fisetin treated female APP^NL-F/NL-F^ mice, but no improvements in memory recall were noted. Although the exact mechanism is unclear, this may suggest a different dosing schedule is needed for fisetin to reach similar efficacy as D+Q treated female APP^NL-F/NL-F^ mice.

The reasons behind the sexually dimorphic responses to the senolytics used in this current study may partially be explained by an earlier onset of disease progression in female APP^NL-F/NL-F^ mice. Although APP^NL-F/NL-F^ mice develop cognitive impairment at 12-18 months of age [18] when hippocampal plaque accumulation is prevalent, we have previously shown females accumulate more plaques compared to males [40]. The present study also highlighted that senescent cell markers (SA-β-gal, p16^Ink4α^) as well as soluble and insoluble Aβ_42_ were elevated in vehicle treated females compared to treatment matched male APP^NL-F/NL-F^ mice. Senolytics clear senescent cells or reverse cellular senescence rather than preventing cells from entering a senescent state. The D+Q treatment may have been efficacious in female APP^NL-F/NL-F^ mice because their disease progression, senescent cell burden, and SASP was more pronounced compared with age-matched C57BL/6 and APP^NL-F/NL-F^ male littermates.

Differences with adiposity and plasma adiponectin may also account for sex-dependent D+Q treatment effects in APP^NL-F/NL-F^ mice. Although D+Q treatment in the male APP^NL-F/NL-F^ mice did not change body weight or WAT accumulation, the treatment drastically reduced circulating levels of adiponectin, which has strong anti-inflammatory properties [41, 42]. The decreased adiponectin in D+Q treated male APP^NL-F/NL-F^ mice could increase SASP in vWAT. However, elevated vWAT, p21^cip1^, and SASP markers in the D+Q treated male mice did not affect plasma cytokines, hippocampal inflammatory senolytic markers, plaque burden, or cognitive function. Although adiponectin levels were unchanged after senolytic intervention in female APP^NL-F/NL-F^ mice, D+Q treatment reduced body weight and WAT depots as well as vWAT and plasma senescent cell and SASP markers.

Metabolic changes also underlie the differential sex responses to senolytic intervention. Besides reduced adiposity, D+Q treated female APP^NL-F/NL-F^ mice had improved energy metabolism potentially contributing to the lower plasma insulin and basal blood glucose levels. Circulating triglyceride levels, associated with a lower AD risk in women [43], were also decreased. Both senolytic treatments increased plasma insulin levels in male APP^NL-F/NL-F^ mice. This suggests impaired glucose utilization, although this was not observed with the ITT or GTT. We suspect that there may be more factors involved in the sex-dependent response of APP^NL-F/NL-F^ mice to the senolytic drugs used in the current study, which warrants further investigation.

APP^NL-F/NL-F^ mice are backcrossed onto a C57BL/6 background strain. Surprisingly, these results contrast with our previous studies utilizing a similar experimental paradigm in C57BL/6 mice [20]. We previously demonstrated that D+Q treatment elevated SASP expression, increased WAT mass, and reduced energy metabolism that decreased cognitive performance in female C57BL/6 mice. The dichotomous results between these two studies may be attributed to factors associated with physiological and pathological aging. Physiological aging involves the natural changes to an organism that occur over time. Pathological aging is not simply accelerated aging. Instead, these are the biological changes associated with disease progression that often cause significant impairment in function and quality of life. APP and the peptides produced by amyloidogenic proteolytic cleavage are known to have detrimental effects on metabolic function both centrally and peripherally [44]. Knock-in of the humanized mutations to APP induces pathological changes starting in early adulthood that deviate metabolic function both centrally and peripherally in APP^NL-F/NL-F^ mice from their C57BL/6 genetic background littermate. Unlike during physiological aging in C57BL/6 mice, the APP knock-in pathologically predisposes mice to amyloid accumulation that drives production of neuroinflammation and induces senescence [45]. When directly comparing vehicle treated mice from our prior study [20], male and female APP^NL-F/NL-F^ mice had worse fed blood glucose and insulin tolerance compared with age-matched C57BL/6 mice. We also noted female APP^NL-F/NL-F^ mice had fewer platform crossings, but D+Q treatment improved this to a similar threshold as sex-matched C57BL/6 vehicle treated mice.

Considering the role of insulin resistance on Aβ_42_ pathological progression, this may be a contributing factor to cognitive decline in this AD model, similar to observations in other amyloidogenic mouse models.

The sexually dimorphic effects of D+Q administration may be due to differences in accumulation of senescent cells when treatment was initiated. Four month old female APP^NL-F/NL-F^ mice had higher expression of several vWAT senescent cell and SASP markers compared to age- and genotype-matched males. This would suggest female APP^NL-F/NL-F^ mice have a faster pathological progression compared to males. At six and 12 months of age, female APP^NL-F/NL-F^ mice have elevated soluble Aβ_42_ compared with age-matched males [46]. While in the present study, we observed larger plaque size in 13 month old female APP^NL-F/NL-F^ mice receiving vehicle treatment. This disparity in amyloid progression between the sexes and its subsequent clearance into plasma may cause the increased peripheral senescence observed in female mice. Despite initiating treatment at the same ages, the faster pathological progression in female APP^NL-F/NL-F^ mice places them at a more advanced disease stage and could account for the sexually dimorphic efficacy of D+Q treatment. Surprisingly, D+Q treatment increased p21^Cip1^ and SASP markers in male APP^NL-F/NL-F^ mice unlike studies in other aging and disease models [38, 47]. This may also indicate that reaching a critical threshold of senescent cell accumulation is necessary, otherwise senolytic intervention may have unintended consequences. While the mechanisms behind this remain to be elucidated, detrimental effects of D+Q administration are not unique to this study. Using the same dose, D+Q treatment worsened male hepatocellular carcinoma in male C57BL/6 mice [48]. Others have proposed that continuous senescent cell removal beginning at young ages does not always activate regenerative processes, but rather causes fibrosis that is detrimental to the healthspan of the organism [49].

Previous research investigating the effects of senolytic compounds in different AD models has yielded varying results. Most of these studies, however, either examined a single sex or pooled data from both sexes making it difficult to identify sex-specific differences. In APP/PS1 mice, females receiving D+Q treatment had reduced plaque burden and improved cognitive performance [15] while fisetin improved cognitive performance in males without affecting amyloid pathology [50]. In the rTg4510 tauopathy model, D+Q treatment reduced neurofibrillary tangles when data from both sexes were pooled [14]. More recently, sexually dimorphic effects of fisetin and D+Q have been observed in rTg4510 mice with cognitive improvements noted in females, but not males [51]. The rTg4510 model has a notable concern when examining therapeutic interventions since overexpression of mutant human tau is not sufficient to cause the phenotypic changes observed in this AD model. D+Q treatment in another tauopathy model, PS19, reduced neurofibrillary tangles and improved cognition, but sex differences were not examined [52]. D+Q treatment administered solely to female 3xTg mice reduced amyloid and tau pathology, with mild memory improvements observed [53]. Despite differences in dosing concentrations and schedules, the disease modifying benefits observed in these prior studies favored: 1) intervention during the primary prevention window, 2) D+Q treatment, and 3) female mice. Which is similar to observations with our current study. This suggests senolytic treatment for AD could be tailored for sex or disease state.

Several study design limitations should be noted. First, was an inability to conduct the *in vivo* assays during a single treatment window. The duration needed to complete the metabolic and cognitive assays required these parameters to be assayed across several months as to avoid confounds due to testing during senotherapeutic dosing weeks. Second, the vehicle used and dosing schedule was modified from prior publications. Senolytic compounds were initially dissolved in canola oil to reduce the concentration of DMSO needed. While canola oil can affect weight and plasma lipids, this would have been consistent across treatment and vehicle control groups. We also used a different Fisetin schedule opting for once per month as opposed to five consecutive days. This was based on our prior work showing efficacy in male C57BL/6 mice. The lack of effects seen on energy metabolism or glucose regulation despite worse circulating concentrations of insulin and adiponectin may be due to fewer senolytic treatments received. However, the trends towards D+Q benefits in females and detriments in males were consistent across the assays. Third, we tested the senolytics in young mice when senescence and SASP markers were elevated compared to C57BL/6 littermates in order to determine their efficacy as a primary interventional strategy. It is possible the compounds may have different therapeutic effects at different stages of disease progression.

Overall, our current study shows that administering D+Q during a primary prevention window reduced central and peripheral markers of senescence and SASP in female APP^NL-F/NL-F^ mice. This led to improvements in adiposity, metabolism, and cognitive performance. Considering women have a greater risk of developing dementia during their lifetime, identifying senotherapeutics appropriate for sex and dementia stage is necessary for personalized AD therapies.

## Abbreviations

AD: Alzheimer’s disease
Aβ: Amyloid beta
APP: amyloid precursor protein
AUC: area under the curve
bw: body weight
BAT: brown adipose tissue
VCO2: carbon dioxide consumption
CIPL: corrected integrated path length
D: Dasatinib
EE: energy expenditure
GTT: glucose tolerance test
GAPDH: glyceraldehyde-3-phosphate dehydrogenase
ITT: insulin tolerance test
i.p.: intraperitoneal
MWM: Morris water maze
VO2: oxygen consumption
Q: Quercetin
RQ: respiratory quotient
SA-β-gal: senescence-associated-β-galactosidase
SASP: senescence associated secretory phenotype
SEM: standard error of the mean
UBE2D2: ubiquitin conjugating enzyme E2 D2
WAT: white adipose tissue
vWAT: visceral white adipose tissue

## Acknowledgements

We would like to thank Melissa Roberts for running the plasma cytokine assay and Stephen Roy for his assistance in editing the manuscript

---

**Online Resource 1.**
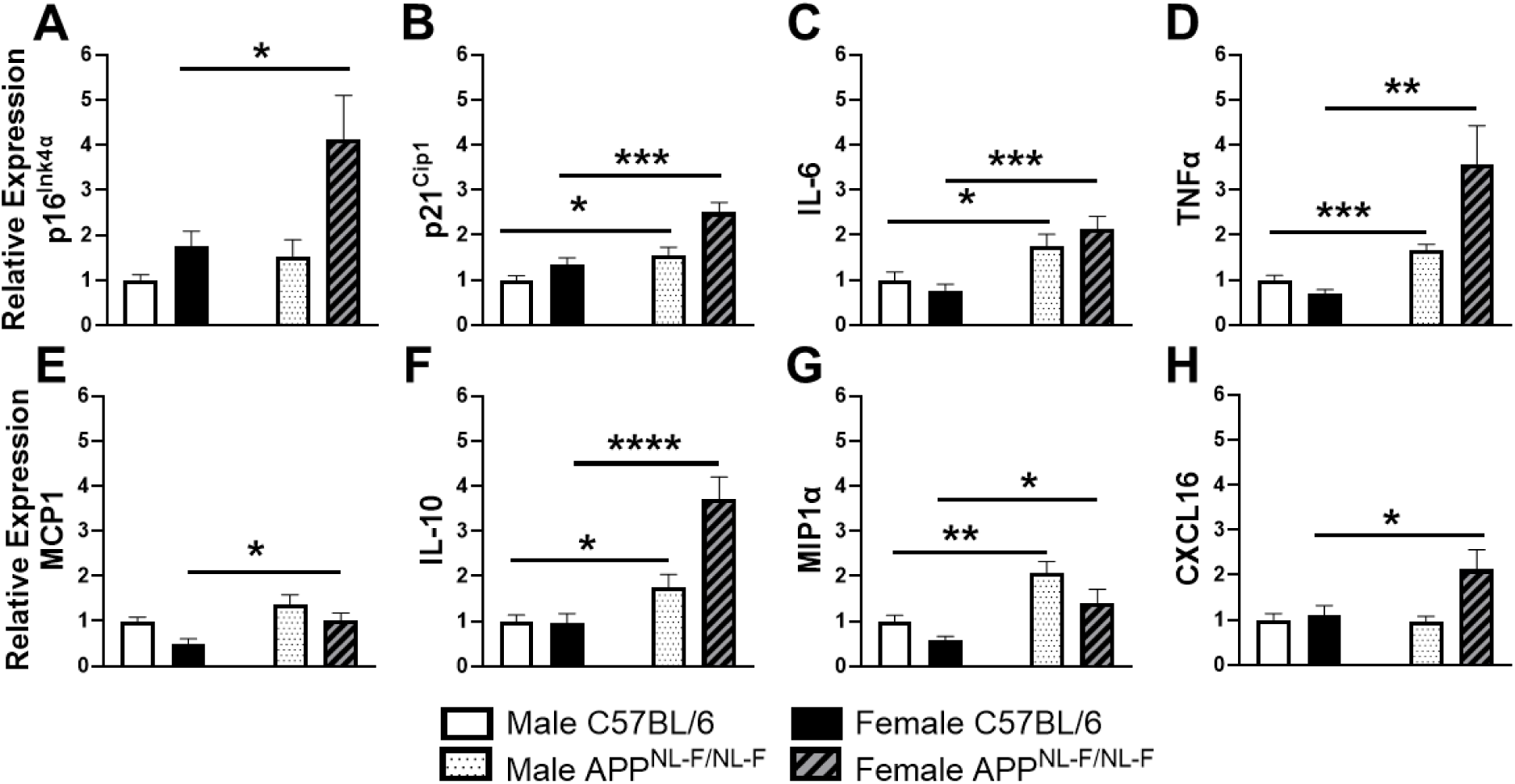
Relative expression of senescent cell and SASP markers in visceral adipose tissue. Gene expression profiles in vWAT (A-H). Data are represented as means ± SEM (n=10-12). *p<0.05, **p<0.01, ***p<0.001, ****p<0.0001 based on a two-tailed Student’s *t* test.

**Online Resource 2.**
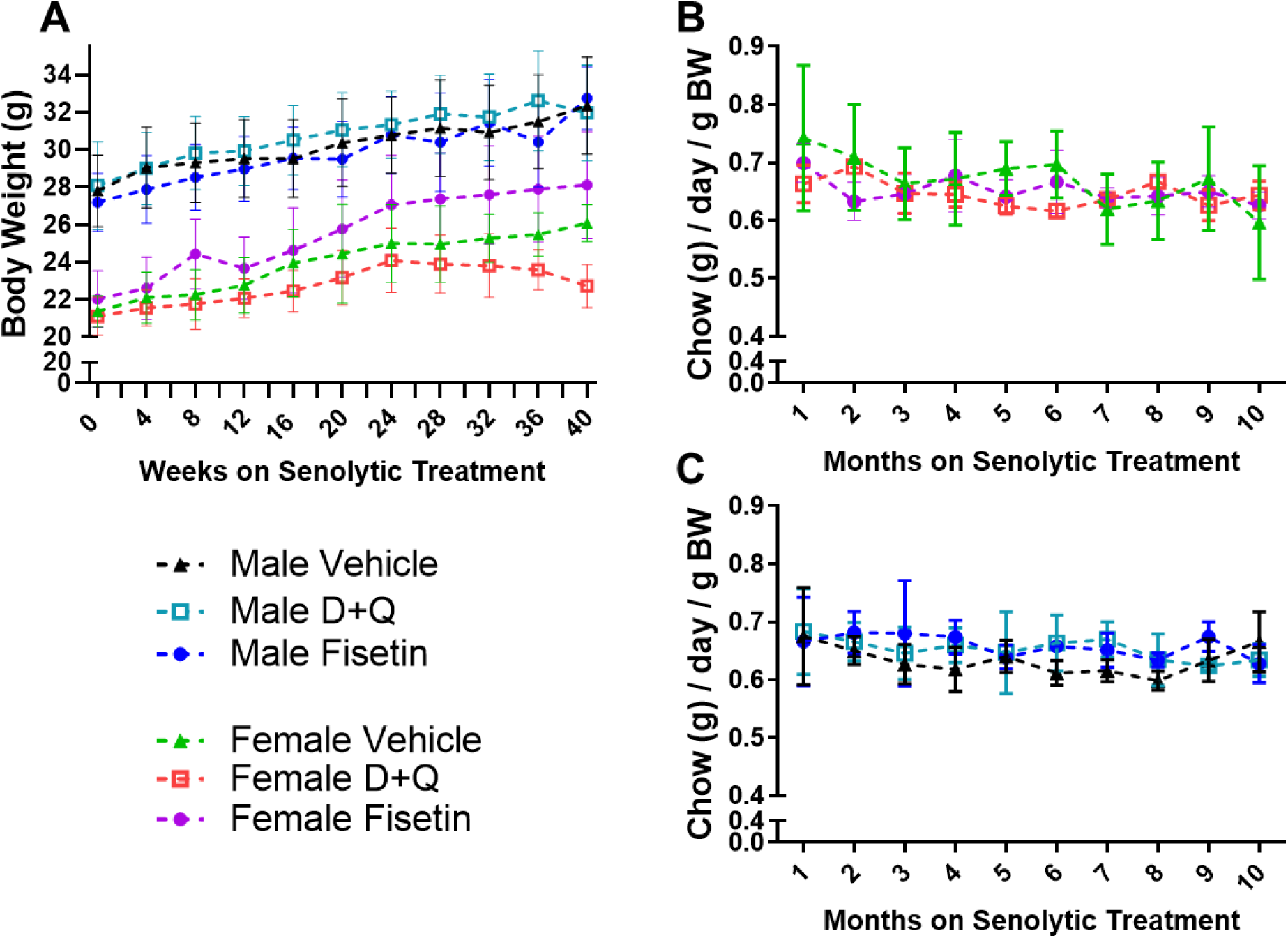
Longitudinal body weight and chow consumption during senolytic treatment. Body weight (BW; A) and chow consumption (B-C) were monitored weekly with the later averaged across treatment months. Time scale denotes start of senolytic treatment when mice were four months of age. Data are represented as means ± SEM (n=5-10).

**Online Resource 3.**
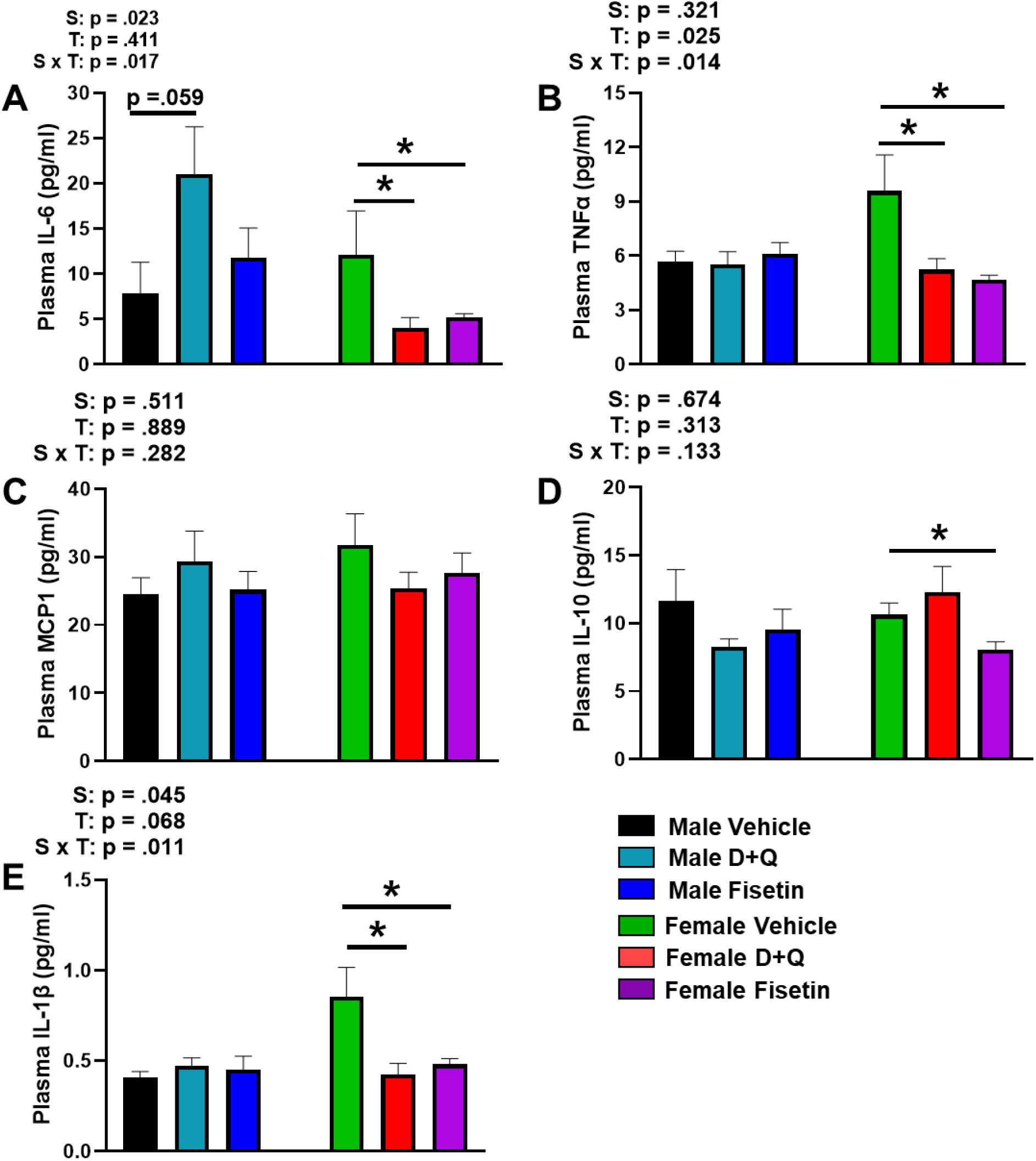
Plasma cytokines after senolytic treatment. Circulating plasma IL-6, TNFα, MCP1, IL-10, and IL-1β (A-E) levels in male and female APP^NL-F/NL-F^ mice. Data are represented as means ± SEM (n=6-10). Results of a two factorial analysis are shown above each bar graph for the Sex (S) and Treatment (T) categorial variables and their interaction (S x T). *p<0.05 based on a two-tailed Student’s *t* test.

